# Ontology-based prediction of cancer driver genes

**DOI:** 10.1101/561480

**Authors:** Sara Althubaiti, Andreas Karwath, Ashraf Dallol, Adeeb Noor, Shadi Salem Alkhayyat, Rolina Alwassia, Katsuhiko Mineta, Takashi Gojobori, Andrew D Beggs, Paul N Schofield, Georgios V Gkoutos, Robert Hoehndorf

## Abstract

Identifying and distinguishing cancer driver genes among thousands of candidate mutations remains a major challenge. Accurate identification of driver genes and driver mutations is critical for advancing cancer research and personalizing treatment based on accurate stratification of patients. Due to inter-tumor genetic heterogeneity, many driver mutations within a gene occur at low frequencies, which make it challenging to distinguish them from non-driver mutations. We have developed a novel method for identifying cancer driver genes. Our approach utilizes multiple complementary types of information, specifically cellular phenotypes, cellular locations, functions, and whole body physiological phenotypes as features. We demonstrate that our method can accurately identify known cancer driver genes and distinguish between their role in different types of cancer. In addition to confirming known driver genes, we identify several novel candidate driver genes. We demonstrate the utility of our method by validating its predictions in nasopharyngeal cancer and colorectal cancer using whole exome and whole genome sequencing.

## Introduction

The natural history of cancer is complex [1] and its genetics highly heterogeneous [2]. Carcinogenesis depends on the accumulation of, and selection for, mutations in genes which are subsequently able to drive, amongst other processes, cell proliferation, immune evasion, genomic instability, and invasiveness. The concept of a cancer driver genes captures the mechanistic requirement for a cell to overcome normal cellular and tissue homeostatic mechanisms and initiate or promote neoplasia and malignancy. The mutations in driver genes are often recessive loss-of-function lesions exemplified by tumor suppressor genes but they can also act dominantly as oncogenes. Determining which mutations or genes act as a bottleneck in the generation of cancer is fraught with problems, as cells carrying one or more driver mutations will also carry a large set of co-selected, “passenger”, mutations which constitute most of the normal somatic mutation-load of the expanded cancer cell but which do not directly generate the neoplastic phenotype [3].

Much effort has gone into developing algorithms to identify driver genes and their mutations, most of which are based on the frequency or pattern of mutations in multiple tumors and their predicted pathogenicity. The goal of identifying cancer drivers may be achieved at the level of gene, protein or pathways, and multiple approaches have been attempted to date [4]. There is no gold standard against which the success of an algorithm can be measured, although the Cancer Gene Census approaches a “gold standard” most closely with an expert-curated dataset of cancer-associated genes and mutations [5].

Investigators have increasingly relied on taking a consensus of multiple methods and, where possible, attempted to experimentally verify driver gene status in cellular or whole organism systems [6]. Many thousands of tumors have now been sequenced in very large-scale studies of multiple cancer types, and several hundred genes and mutations have been identified as “drivers” – with varying support from experimental and genetic studies [6]. These methods, however, do not work well for low-intermediate and rare driver genes which may bear up to 20% of driver mutations [7], and the identification of drivers in specific cancers and sub-types of tumor remains difficult, often because of small numbers of tumors available.

An alternative strategy to sequence-based ratiometric type mutation frequency based approaches is to identify a “fingerprint” for cancer driver genes from a range of biological and molecular data and to use this as part of a classifier that can filter sequence information. For example, it is possible to utilize gene annotations and biological properties of known driver genes in a machine learning approach and identify some novel driver gene candidates [8]. Here, we report a novel method in which we use a combination of direct functional evidence, obtained through cell growth assays after introducing a specific mutation into cells, and related functional characteristics available from basic biological experiments and recorded in model organism databases and databases characterizing protein functions to build a classifier that determined whether a gene may likely become a driver gene.

Public databases contain large volumes of information that relates genes or variants to phenotypes (either on the cellular or whole body organism level), the specific biological processes and molecular functions they can be involved in, or the cellular locations at which a gene product is active. Phenotypes are systematically collected in the context of genotype–phenotype relations, both from human clinical information [9], from model organism experiments [10], and for cell models in cellular phenotype databases [11]. Information about gene functions is collected in databases such as Uniprot [12] as well as several model organism databases.

In these databases, ontologies are used for characterizing phenotypes, gene functions and cellular locations. Over the past decades, a tightly integrated system of ontologies has been developed that interlinks the knowledge about basic biological phenomena through the use of logical axioms [13]. Exploring the information in this system of ontologies can enable novel types of analysis [14] and the background knowledge in the ontologies has the potential to significantly improve biomedical data analysis [15].

We have developed a method that uses biological background knowledge about the relation between genes or variants and their phenotypes, either on the cellular or whole body organism level, as well as gene functions and cellular locations, to predict driver genes and mutations. Our approach relies on neuro-symbolic deep learning to systematically encode for background knowledge about basic biological processes and phenomena. Specifically, we generate “embeddings” for gene functions and gene–phenotypes associations and use a deep artificial neural network trained on known driver genes – and the knowledge about how they relate to functions and phenotypes – to discover new cancer drivers. We demonstrate that our method can predict novel driver genes by analyzing three cohorts of different cancers, and we demonstrate that the predicted driver genes have a significantly higher somatic mutation frequency, are significantly more likely to be functionally related to known drivers, and have a significantly higher rate of pathogenic somatic variants. We make our method and prediction results freely available from https://github.com/bio-ontology-research-group/predCAN.

## Results

### Representation learning

Our aim is to utilize information about the functions and phenotypes associated with genes to identify cancer driver genes and, subsequently, driver mutations. This information is represented using biomedical ontologies, and these ontologies also contain a substantial amount of background information about the relations between biological functions, processes, and phenotypes in the form of logical axioms and natural language definitions [14]. The information in ontologies is utilized by human experts to understand and interpret the implications of an association with a class in an ontology, and a comprehensive interpretation of these associations relies on comprehension and utilization of biological background knowledge.

We use three types of information associated with genes: cellular phenotypes observed in large-scale microscopy studies and recorded using the Cellular Microscopy Phenotype Ontology (CMPO) [16]; gene functions and cellular locations recorded by Uniprot [12] and encoded using the Gene Ontology (GO) [17]; and phenotypes of knockout mouse models provided by the Mouse Genome Informatics (MGI) database [10] and encoded using the Mammalian Phenotype Ontology (MP) [18].

Each of these ontologies contains logical axioms that define and restrict the classes and provide background knowledge about the domain of functions, cellular phenotypes, or physiological phenotypes [18]. Figure 1 shows an example of how the ontologies overlap in their content and how classes in phenotype ontologies are defined or constrained using classes from other ontologies.

**Fig 1.**
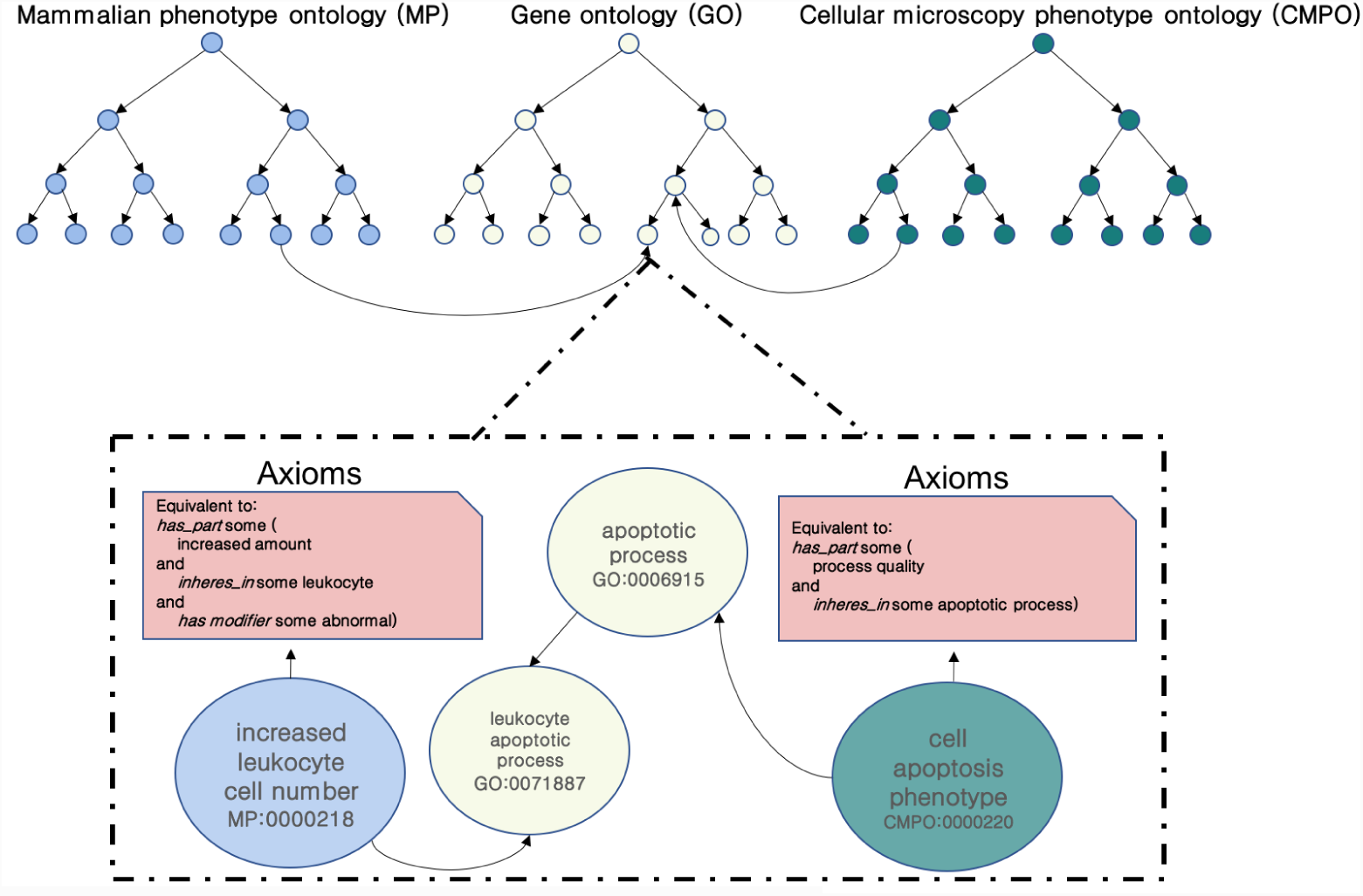
Interlinked knowledge between ontologies.

We developed a method that combines the associations of genes and ontology classes with the background information in ontologies (both the formal logical content and the informal natural language components) in a single feature vectors that represents each gene in our dataset. For this purpose, we utilize neuro-symbolic feature learning on ontologies in which machine learning models are combined with deductive inference on formally represented knowledge [19]; as a result, we obtain an embedding – a function from an ontology and its associated entities into an *n*-dimensional vector space – which generates feature vectors that encode for known associations of genes and their functions or phenotypes as well as the ontologies’ background knowledge.

Because different genes are covered differently in the databases we use, we generate five different representations for gene. The first three representations are based on annotating the genes using the ontologies that we used one at a time, combining the ontology-based annotations and the ontology axioms within a single representation so that the background knowledge within the ontologies becomes accessible. However, we can only generate the feature vectors if there are ontology-based annotations for a gene and therefore we obtain a different number of feature vectors when utilizing different ontologies. To determine if we can predict better, we combine the embeddings generated from each gene if all three features are available and evaluate the performance on the intersection of genes for which features in all three ontologies can be generated. Finally, we determined if it is possible to “impute” missing information by utilizing ontology axioms. For this purpose, we first merge the three ontology (CMPO, GO, MP) into a single ontology model and generate the embeddings for each gene which is annotated to information in at least one of the three ontologies we use. We use the generated feature vectors as input to a deep neural network that we train to predict cancer driver genes and distinguish between 20 cancer types for which a gene can be considered a driver.

### Biological features and background knowledge predict cancer drivers

We first test how well each type of feature predicts cancer driver genes separately. For this purpose, we construct a machine-learning model to classify genes into driver genes for particular cancer types; we distinguish between 20 different types of cancer (see Supplementary Table 1) taken from the IntoGen database [20]. The machine learning model we use is illustrated in Supplementary Figure 1. We evaluate the results using 10-fold cross-validation and performance results are summarized in Table 1. We then combine the features and the ontologies that contain background knowledge about the respective domains; when adding ontologies containing background knowledge about the cellular phenotypes and functions of genes we achieve a significant improvement of prediction results compared to predicting based on individual features (see Table 1, Figure 2 for ROC curves and Figure 3 for the precision-recall curves). As our method can accurately predict cancer driver genes, we apply our model to all human genes (i.e., all negatives in our dataset) and predict 112 novel candidate driver genes for 20 different cancer types (Supplementary Tables 2 and 3).

**Table 1.** ANN performance in identification driver/non-driver genes of different cancer types. We first determine the performance using each ontology-based feature separately (using each ontology individually as background knowledge), and then determine the performance only on genes for which we have all three ontology-based features (Intersection); in the “Intersection” case, embedding vectors are concatenated. Finally, we determine the performance on genes for which we have at least one feature (Union); in the latter case, ontologies are merged.

| Experiments | F-score | AUC | Number of annotated genes |
| --- | --- | --- | --- |
| CMPO | 78.95% | 72.96% | 13,116 |
| GO | 84.23% | 78.81% | 17,591 |
| MP | 85.39% | 79.51% | 7,884 |
| CMPO+GO+MP (Intersection) | 91.20% | 89.73% | 7,884 |
| CMPO+GO+MP (Union) | 92.57% | 94.28% | 20,352 |

**Fig 2.**
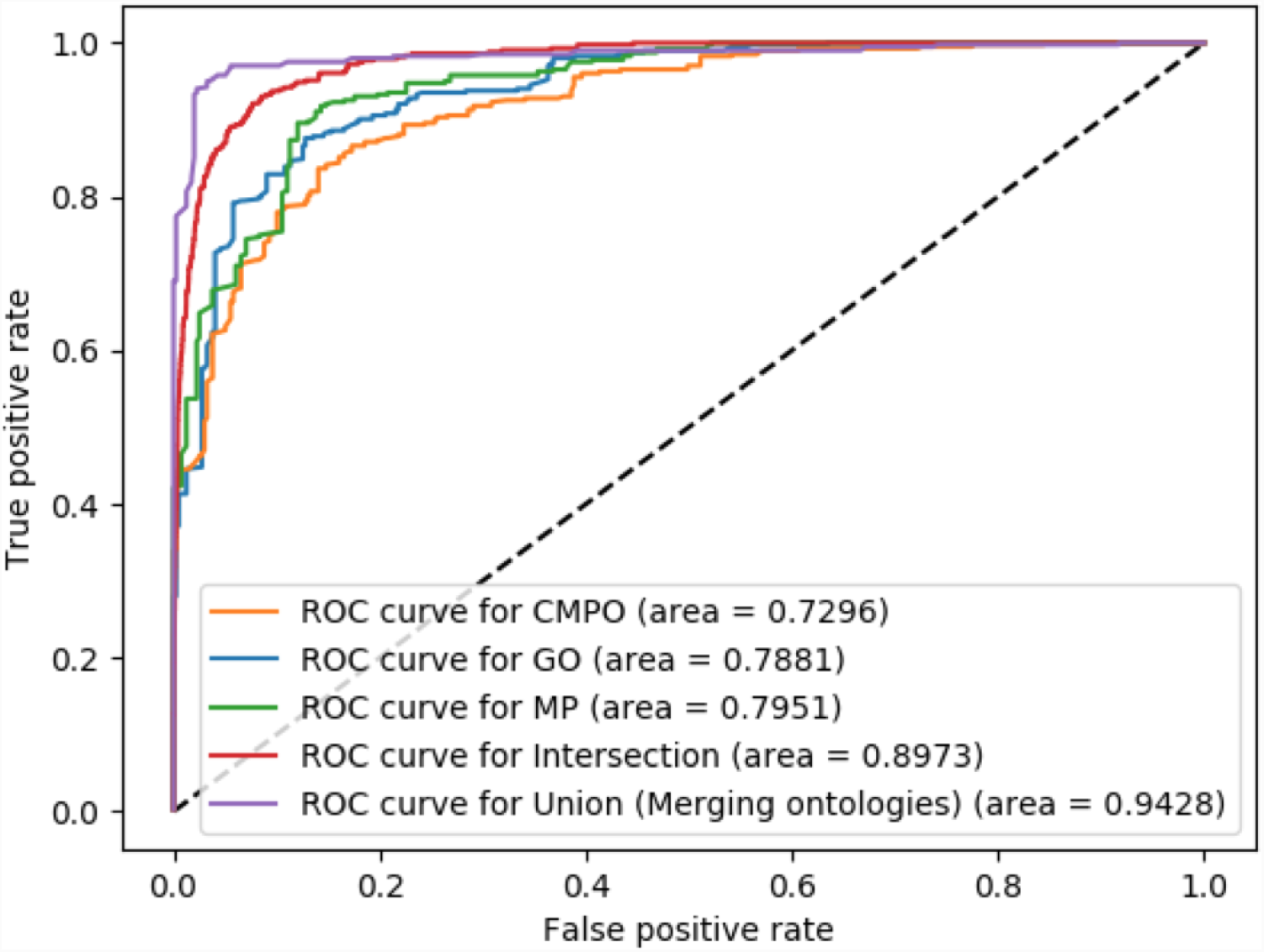
ROC curves and AUC values for the prediction of driver genes based on different ontology associations.

**Fig 3.**
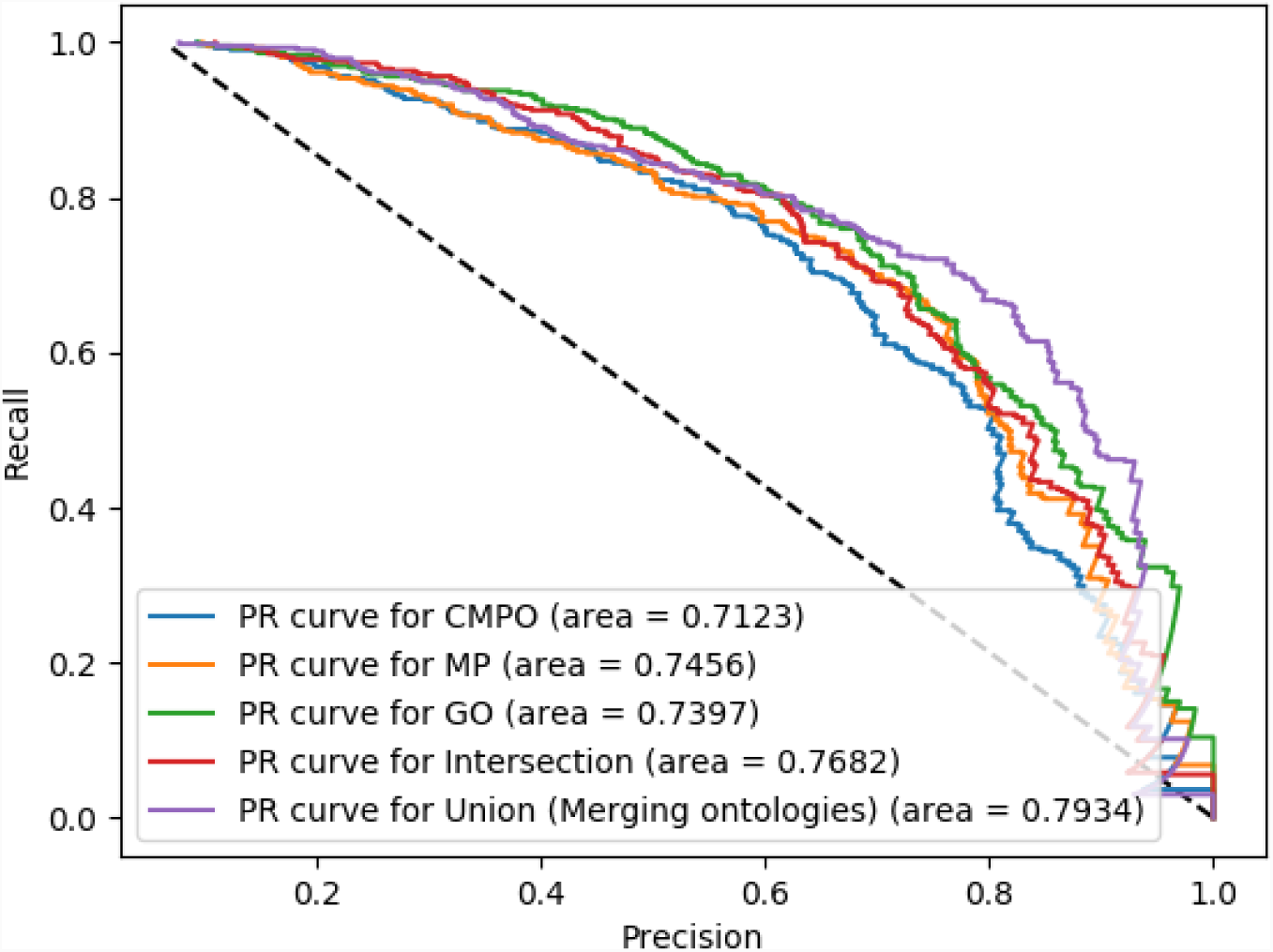
Precision-recall curves for the prediction of driver genes based on different ontology associations.

**Fig 4.**
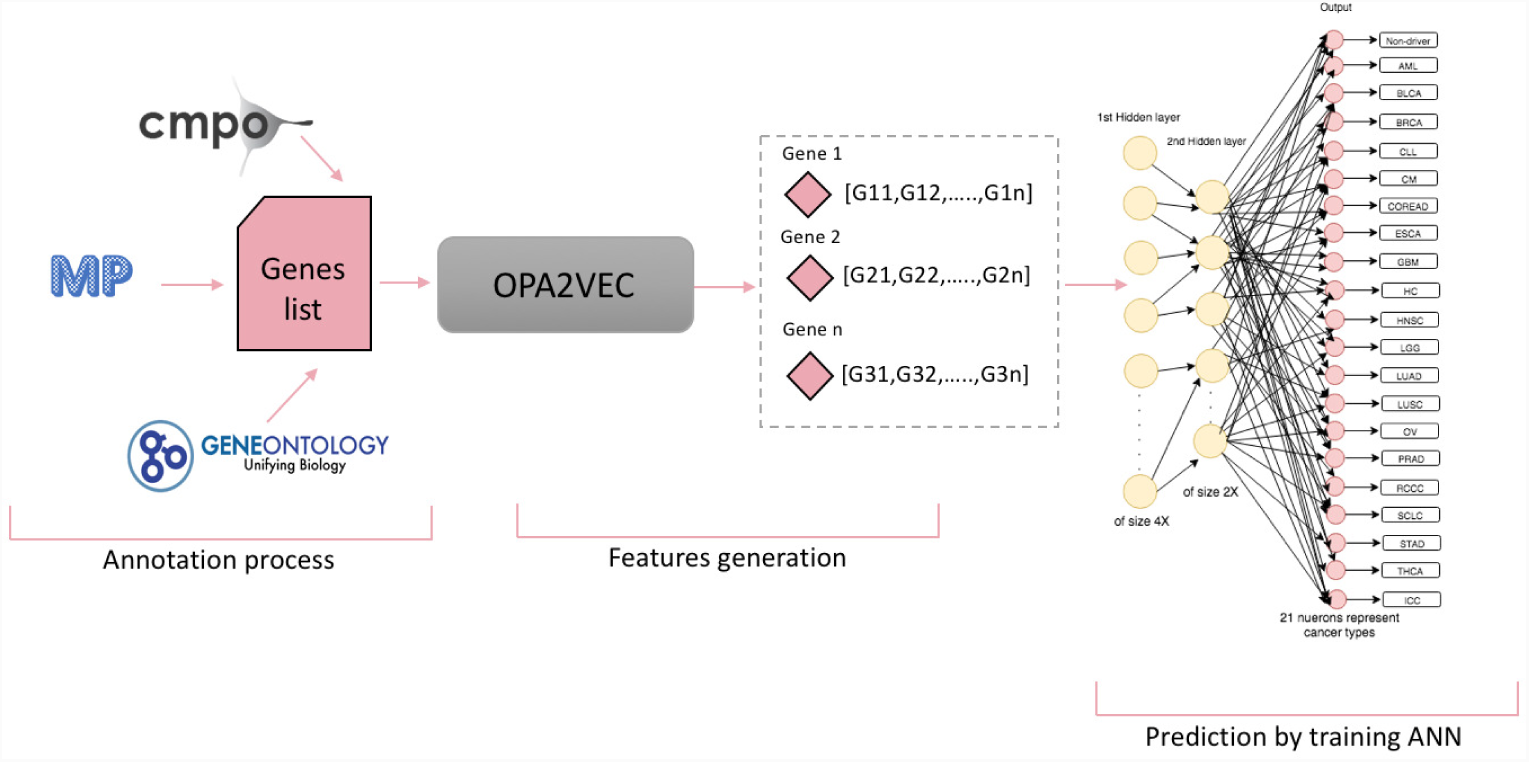
predCAN workflow. Starting with annotation the genes from CPD with CMPO, GO and MP; and generates vectors with OPA2Vec. The prediction process done by training ANN in order to which cancer type each gene belong to discover new driver genes.

### Predicted cancer driver genes are hypermutated in cancer exomes

Driver genes will generally accumulate more mutations than other genes in cancer tissue [21]. We use the complete list of genes in the IntoGen database and separate them into three groups: genes listed as confirmed cancer drivers, genes predicted as cancer drivers for one or more tumor types by our method, and genes not predicted or known as cancer driver genes. We use the recorded somatic mutations in the IntoGen database to determine the somatic mutation frequency for each human gene (i.e., the number of mutations divided by the length of the gene). We find that candidate driver genes predicted by our method have a significantly higher somatic mutation frequency than non-driver genes (*p* = 0.560 × 10^-12^, one-tail t-test).

We also evaluate our predictions on a set of 26 tumor samples for nasopharyngeal cancer obtained from patients at King Abdulaziz University Hospital in Jeddah, Saudi Arabia, for which we have performed whole exome sequencing. Nasopharyngeal cancer is a type of squamous cell carcinoma of the head and neck for which we identify seven novel candidate driver genes (see Supplementary Table 3). In this set of samples, the predicted driver genes have a significantly higher somatic mutation frequency compared to non-driver genes (*p* = 0.115 × 10^-5^, one-tail t-test).

Moreover, we applied the same analysis to 114 tumor/normal samples for colorectal adenocarcinoma obtained from the University of Birmingham Hospital in UK. We identified 12 candidate driver genes for colorectal adenocarcinoma. Among the 114 samples, the predicted driver genes have a significantly higher somatic mutation frequency compared to non-driver genes (*p* = 0.204 × 10^-3^, one-tail t-test).

### Predicted cancer driver genes are functionally related to known cancer drivers

Cancer drivers are known to form functional modules on interaction networks [22]. We use the STRING interaction network [23] to determine whether the genes we predict are functionally related to known driver genes. STRING contains several types of interaction between genes and proteins, including physical interactions, genetic interactions, co-location, and co-expression.

We find that 24 out of the 112 candidate driver genes have a direct interaction with a known driver gene within their respective cancer type. In a random distribution (see Methods), on average six genes are connected to a known driver gene, demonstrating that our predicted drivers are significantly more likely to be functionally related to a known driver gene (*p* = 0.731 × 10^-7^, one-tail t-test).

### Predicted driver genes are enriched for pathogenic variants

Our method can also be used to detect driver variants in tumor samples. We use seven different methods for predicting deleteriousness of variants and compare the pathogenicity scores of variants in candidate driver genes to known and non-driver genes in nasopharyngeal carcinoma exomes. Variants in the seven predicted driver genes for head and neck squamous cell carcinoma generally are scored more pathogenic than variants non-driver genes by most prediction tools (SIFT: *p* = 0.324 × 10^-5^, PolyPhen-2: *p* = 0.098, MutationAssessor: *p* = 0.378 × 10^-6^, MutationTaster: *p* = 0.372 × 10^-10^, CADD: *p* = 0.818, VEST3: *p* = 0.169 × 10^-12^ and FATHMM: *p* = 0.912 × 10^-14^; Mann-Whitney U test).

We apply the same test on the cohort of colorectal adenocarcinoma samples. The set of variants in the predicted driver genes for colorectal adenocarcinoma (see Supplementary Table 5) are scored as significantly more pathogenic than variants in non-driver genes by most pathogenicity prediction methods (SIFT: *p* = 0.631 × 10^-6^, PolyPhen-2: *p* = 0.725 × 10^-8^, MutationAssessor: *p* = 0.405 × 10^-2^, MutationTaster: *p* = 0.917, CADD: *p* = 0.811, VEST3: *p* = 0.040 and FATHMM: *p* = 0.648 × 10^-8^; Mann-Whitney U test).

We can also use these methods to suggest candidate driver mutations in the cohorts as well as individual patients. Supplementary Tables 4 and 5 lists all rare variants in predicted driver genes in the predicted driver genes for nasopharyngeal carcinoma and colorectal cancer. For example, *TGM3* has previously been studied as candidate driver in carcinomas of the head and neck [24], and we identify `rs200294064` in *TGM3* as pathogenic candidate driver mutation.

## Discussion

Our method can predict cancer driver genes using only public background knowledge about gene functions, cellular and organism phenotypes. The key novelty of our algorithm is the ability to encode biological background knowledge in ontologies [15], and therefore exploit inter-ontology links in the form of axioms; the predictive performance of our method is best when combining different – yet related – ontologies. While many of the axioms that relate classes in different ontologies have been created to support ontology development and maintenance, we show that ontologies in the biomedical domain now form a comprehensive web of biological background knowledge that can – when exploited through appropriate learning algorithms – significantly improve the interpretation and analysis of data. A crucial role for this type of connection between different ontologies is the Relation Ontology [25] which is central to ontologies that make up the OBO Foundry [13].

Through the application of our method we identify novel driver genes by systematically analyzing public knowledge on multiple levels of granularity. We use sequencing data – both public and patient-derived – as validation. This strategy is the opposite of most computational and experimental approaches in which molecular data is used as feature and additional functional evidence collected after predictions [6,26]. Our approach can also identify consensus cancer driver genes as we found 8 out of the 112 have been listed in the current COSMIC of confirmed consensus cancer driver genes [27] (see Supplementary Table 6). Moreover, we found 12 out of the 26 rare variants within the predicted driver genes for both nasopharyngeal and colorectal cohorts mentioned in International Cancer Genome Consortium (ICGC) data [28] (see Supplementary Table 7). One of the key novelties in our approach is the use of phenotype data – both on the cellular and whole organism levels of granularity – to characterize cancer driver genes.

## Materials and methods

### Data sources and ontologies

We performed all our experiments on a set of driver/non-driver genes from the Mutational Cancer Drivers Database (intOGen) [20] and Cellular Phenotype Database (CPD) [11] downloaded on 18 Feb 2018. They contain a total of 60,279 genes: 24,475 of them are human protein-coding genes and the remaining 35,804 represent different types of RNA molecules and pseudogenes. Some of the genes are identified as being driver genes in one of the 28 cancer types in intOGen.

Furthermore, we used cellular phenotype annotations of genes provided by the Cellular Phenotype Database (CPD) [11], downloaded on 18 Feb 2018, and we used function annotations provided by the Gene Ontology (GO) website [17] downloaded on 14 Jul 2018. We further used phenotype annotations observed in mutant mouse models, downloaded from the Mouse Genome Informatics (MGI) database [10] on 14 Jul 2018.

We mapped all gene identifiers to Entrez gene identifiers and used the mapping file between Ensembl gene identifiers and Entrez genes provided by the Cellular Phenotype Database (CPD) [11]; we converted UniProt identifiers of proteins in the GO annotations to Entrez gene identifiers using the UniProt mapping tool provided on the UniProt website [12]; and for assigning phenotypes to genes we use mouse orthologs of human genes and assign the phenotypes associated with loss-of-function mutations in mice in the MGI database [10] to the human genes.

We started with 20,352 human genes of which 13,116 are annotated using the Cellular Microscopy Phenotype Ontology (CMPO) [16], 17,591 are annotated using the Gene ontology (GO) [17], and 7,884 are annotated with the Mammalian Phenotype Ontology (MP) [18].

### Generating of ontology annotation-based features

We used the OPA2Vec approach [19] to generate “embeddings” representing genes based on the different ontologies and the annotations of the genes with the ontologies. An ontology embedding is a function that projects entities in an ontology or annotated with an ontology into an *n*-dimensional real-valued space.

We generated ontology embedding for each gene using the following parameter settings of OPA2Vec: embedding size 100, window size 5, minimum count 5, and using a skip-gram model.

### Supervised training

We investigated the performance of a machine learning based classification algorithm in identifying new driver genes. In our experiments, we used known driver genes within 20 different cancer types as the positive set, and randomly selected an equal number of the non-driver genes as the negative set.

For the training and testing, we performed stratified 10-fold cross-validation. We performed a limited grid search for optimal sets of hyperparameters of the neural network using the Hyperas system [29]. We used a Rectified Linear Unit as an activation function [30] for the hidden layers and a sigmoid function as the activation function for the output layer; we used cross entropy as loss function in training, and Rmsprop [31] to optimize the neural networks parameters in training.

### Sample preparation and sequencing

For nasopharyngeal cancer samples: Tissue samples were taken from nasopharyngeal cancer lesions and embedded in formalin at King Abdulaziz University Hospital in Jeddah, KSA. DNA was extracted from the tissue using Qiagen QIAamp FFPE Tissue DNA extraction kit (56404) following the manufacturer’s instruction. To capture the exomes and to prepare the sequencing libraries, a TruSeq exome kit (Illumina) was used. The indexed libraries were pooled (7 samples per lane), and 7 lanes in total were used for paired-end sequencing (150bp) on a HiSeq4000 (Illumina). The average sequencing depth is 237x.

For colorectal carcinoma samples: Colorectal cancer tissue and paired germline samples were obtained at the time of surgery and snap frozen in liquid nitrogen until required. For whole genome sequencing, approximately 1-3*μ*g of DNA was sheared and prepared using an Illumina TruSeq PCR-free library preparation kit. Samples were indexed and each sample was pooled across 4 lanes of an Illumina HiSeq 4000 using v4 sequencing chemistry, generating approximately 150Gb of sequencing data per sample using a 125bp PE sequencing strategy at an average depth of 53.7x.

### Filtration for variant calls

For nasopharyngeal cancer samples we used GATK with Mutect2 [32] for sample preparation and generating somatic short variant calling files following tumor-only mode. For colorectal carcinoma, Strelka [33] was used with tumor/normal pairs to detect somatic mutations. We filter all variants that do not lie within a coding region in both cohorts.

We annotate the resulting VCF files with Annovar [34] to extract variant-related information such as the gene(s) in which a variant lies and the start and end position of the gene, and the different pathogenicity scores.

### Detection of network modules

To determine a neutral distribution of the number of connections between cancer driver genes within STRING empirically, we repeatedly sample 112 random nodes from the STRING interaction network [23] and calculate the number of genes which are connected to known cancer driver genes. We repeat this process 10,000 times.

## Acknowledgments

RH was supported by funding from King Abdullah University of Science and Technology (KAUST) Office of Sponsored Research (OSR) under Award No. URF/1/3454-01-01 and FCC/1/1976-08-01. SA, PNS, GVG, and RH were supported by funding from King Abdullah University of Science and Technology (KAUST) Office of Sponsored Research (OSR) under Award No. FCS/1/3657-02-01.

ADB acknowledges funding from the Wellcome Trust (102732/Z/13/Z), Cancer Research UK (C31641/A23923) and the Medical Research Council (MR/M016587/1).

GVG acknowledges support from H2020-EINFRA (731075) and the National Science Foundation (IOS:1340112) as well as support from the NIHR Birmingham ECMC, NIHR Birmingham SRMRC and the NIHR Birmingham Biomedical Research Centre and the MRC HDR UK. The views expressed in this publication are those of the authors and not necessarily those of the NHS, the National Institute for Health Research, the Medical Research Council or the Department of Health.

## Ethics

This work has been reviewed and approved by the Research Ethics Committee of King Abdulaziz University on 29 March 2017 under reference number 116-17, and the Institutional Bioethics Committee at King Abdullah University of Science and Technology on 15 May 2017 under reference number 17IBEC07-Hoehndorf. Ethical approval for work on colorectal cancer samples was obtained via the University of Birmingham Human Biomaterials Resource Centre Biobanking ethics (Ref 09/H1010/75).

## Data access and availability

All data except human genomic data is available from https://github.com/bio-ontology-research-group/predCAN.

Genomic data for nasopharyngeal and colorectal cancer samples cannot be shared publicly due to ethical and legal restrictions. Requests for access can be sent to the University of Birmingham Human Biomaterials Resource Centre Biobanking and the Research Ethics Committee of King Abdulaziz University in Jeddah which will decide on access for researchers who meet the criteria for access to confidential data.

## References

1. Hanahan D, Weinberg RA. Hallmarks of cancer: the next generation. Cell. 2011;144(5):646–74. doi:10.1016/j.cell.2011.02.013.

2. Lawrence MS, Stojanov P, Polak P, Kryukov GV, Cibulskis K, Sivachenko A, et al. Mutational heterogeneity in cancer and the search for new cancer-associated genes. Nature. 2013;499(7457):214–218. doi:10.1038/nature12213.

3. Garraway L, Lander E. Lessons from the Cancer Genome. Cell. 2013;153(1):17–37. doi:https://doi.org/10.1016/j.cell.2013.03.002.

4. Tokheim CJ, Papadopoulos N, Kinzler KW, Vogelstein B, Karchin R. Evaluating the evaluation of cancer driver genes. Proceedings of the National Academy of Sciences. 2016;113(50):14330–14335.

5. Sondka Z, Bamford S, Cole CG, Ward SA, Dunham I, Forbes SA. The COSMIC Cancer Gene Census: describing genetic dysfunction across all human cancers. Nat Rev Cancer. 2018;18(11):696–705. doi:10.1038/s41568-018-0060-1.

6. Bailey MH, Tokheim C, Porta-Pardo E, Sengupta S, Bertrand D, Weerasinghe A, et al. Comprehensive Characterization of Cancer Driver Genes and Mutations. Cell. 2018;173(2):371–385.e18. doi:https://doi.org/10.1016/j.cell.2018.02.060.

7. Lawrence MS, Stojanov P, Mermel CH, Robinson JT, Garraway LA, Golub TR, et al. Discovery and saturation analysis of cancer genes across 21 tumour types. Nature. 2014;505:495. doi:10.1038/nature12912.

8. Chen Y, Hao J, Jiang W, He T, Zhang X, Jiang T, et al. Identifying potential cancer driver genes by genomic data integration. Scientific Reports. 2013;3:3538. doi:10.1038/srep03538.

9. Landrum MJ, Lee JM, Riley GR, Jang W, Rubinstein WS, Church DM, et al. ClinVar: public archive of relationships among sequence variation and human phenotype. Nucleic acids research. 2013;42(D1):D980–D985.

10. Eppig JT, Blake JA, Bult CJ, Kadin JA, Richardson JE, Group MGD. The Mouse Genome Database (MGD): facilitating mouse as a model for human biology and disease. Nucleic acids research. 2014;43(D1):D726–D736.

11. Kirsanova C, Brazma A, Rustici G, Sarkans U. Cellular phenotype database: a repository for systems microscopy data. Bioinformatics. 2015;31(16):2736–2740.

12. Consortium U. UniProt: a hub for protein information. Nucleic acids research. 2014;43(D1):D204–D212.

13. Smith B, Ashburner M, Rosse C, Bard J, Bug W, Ceusters W, et al. The OBO Foundry: coordinated evolution of ontologies to support biomedical data integration. Nat Biotech. 2007;25(11):1251–1255.

14. Hoehndorf R, Schofield PN, Gkoutos GV. The role of ontologies in biological and biomedical research: a functional perspective. Briefings in Bioinformatics. 2015;doi:10.1093/bib/bbv011.

15. Smaili FZ, Gao X, Hoehndorf R. Formal axioms in biomedical ontologies improve analysis and interpretation of associated data. bioRxiv. 2019;doi:10.1101/536649.

16. Jupp S, Malone J, Burdett T, Heriche JK, Williams E, Ellenberg J, et al. The cellular microscopy phenotype ontology. Journal of biomedical semantics. 2016;7(1):28.

17. Ashburner M, Ball CA, Blake JA, Botstein D, Butler H, Cherry JM, et al. Gene Ontology: tool for the unification of biology. Nature genetics. 2000;25(1):25.

18. Smith CL, Goldsmith CAW, Eppig JT. The Mammalian Phenotype Ontology as a tool for annotating, analyzing and comparing phenotypic information. Genome biology. 2005;6(1):R7.

19. Smaili FZ, Gao X, Hoehndorf R. OPA2Vec: combining formal and informal content of biomedical ontologies to improve similarity-based prediction. arXiv preprint arXiv:180410922. 2018;.

20. Perez-Llamas C, Gundem G, Lopez-Bigas N. Integrative cancer genomics (IntOGen) in Biomart. Database. 2011;2011.

21. Stratton MR, Campbell PJ, Futreal PA. The cancer genome. Nature. 2009;458(7239):719.

22. Hofree M, Shen JP, Carter H, Gross A, Ideker T. Network-based stratification of tumor mutations. Nature methods. 2013;10(11):1108.

23. Szklarczyk D, Franceschini A, Wyder S, Forslund K, Heller D, Huerta-Cepas J, et al. STRING v10: protein–protein interaction networks, integrated over the tree of life. Nucleic acids research. 2014;43(D1):D447–D452.

24. Wu X, Cao W, Wang X, Zhang J, Lv Z, Qin X, et al. TGM3, a candidate tumor suppressor gene, contributes to human head and neck cancer. Molecular Cancer. 2013;12(1):151. doi:10.1186/1476-4598-12-151.

25. Smith B, Ceusters W, Klagges B, Köhler J, Kumar A, Lomax J, et al. Relations in biomedical ontologies. Genome Biol. 2005;6(5):R46. doi:http://dx.doi.org/10.1186/gb-2005-6-5-r46.

26. Sjöblom T, Jones S, Wood LD, Parsons DW, Lin J, Barber TD, et al. The Consensus Coding Sequences of Human Breast and Colorectal Cancers. Science. 2006;314(5797):268–274. doi:10.1126/science.1133427.

27. Forbes SA, Bindal N, Bamford S, Cole C, Kok CY, Beare D, et al. COSMIC: mining complete cancer genomes in the Catalogue of Somatic Mutations in Cancer. Nucleic acids research. 2010;39(Suppl 1):D945–D950.

28. Zhang J, Baran J, Cros A, Guberman JM, Haider S, Hsu J, et al. International Cancer Genome Consortium Data Portal—a one-stop shop for cancer genomics data. Database. 2011; 2011. doi:10.1093/database/bar026.

29. Pumperla M. Keras + Hyperopt: A very simple wrapper for convenient hyperparameter optimization; 2016. https://github.com/maxpumperla/hyperas.

30. Nair V, Hinton GE. Rectified linear units improve restricted boltzmann machines. In: Proceedings of the 27th international conference on machine learning (ICML-10); 2010. p. 807–814.

31. Hinton G, Srivastava N, Swersky K. Lecture 6a overview of mini-batch gradient descent (2012). Coursera Lecture slides https://classcourseraorg/neuralnets-2012-001/lecture. 2012;.

32. McKenna A, Hanna M, Banks E, Sivachenko A, Cibulskis K, Kernytsky A, et al. The Genome Analysis Toolkit: a MapReduce framework for analyzing next-generation DNA sequencing data. Genome research. 2010;.

33. Saunders CT, Becq J, Murray LJ, Cheetham RK, Swamy S, Wong WSW. Strelka: accurate somatic small-variant calling from sequenced tumor–normal sample pairs. Bioinformatics. 2012;28(14):1811–1817. doi:10.1093/bioinformatics/bts271.

34. Wang K, Li M, Hakonarson H. ANNOVAR: functional annotation of genetic variants from high-throughput sequencing data. Nucleic acids research. 2010;38(16):e164–e164.

