## Supplementary File 1 for "Ontology-based prediction of cancer driver genes"

Table 1: 20 different cancer types have been used

| Corresponding Class | Cancer Type | Number of genes |
|---|---|---|
| 0 | Not identified | 19,893 |
| 1 | Acute myeloid leukemia (AML) | 32 |
| 2 | Bladder carcinoma (BLCA) | 156 |
| 3 | Breast carcinoma (BRCA) | 184 |
| 4 | Chronic lymphocytic leukemia (CLL) | 38 |
| 5 | Cutaneous melanoma (CM) | 250 |
| 6 | Colorectal adenocarcinoma (COREAD) | 95 |
| 7 | Esophageal carcinoma (ESCA) | 98 |
| 8 | Glioblastoma multiforme (GBM) | 75 |
| 9 | Hepatocarcinoma (HC) | 30 |
| 10 | Head and neck squamous cell carcinoma (HNSC) | 167 |
| 11 | Lower grade glioma (LGG) | 50 |
| 12 | Lung adenocarcinoma (LUAD) | 181 |
| 13 | Lung squamous cell carcinoma (LUSC) | 147 |
| 14 | Ovarian Carcinoma (OV) | 83 |
| 15 | Prostate adenocarcinoma (PRAD) | 88 |
| 16 | Renal clear cell carcinoma (RCCC) | 105 |
| 17 | Small cell lung carcinoma (SCLC) | 61 |
| 18 | Stomach adenocarcinoma (STAD) | 175 |
| 19 | Thyroid carcinoma (THCA) | 32 |
| 20 | Uterine corpus endometrioid carcinoma (ICC) | 149 |

Table 2: Number of predicted driver genes within each cancer type

| Cancer Type | Number of predicted genes |
|---|---|
| AML | 2 |
| BLCA | 4 |
| BRCA | 14 |
| CLL | 2 |
| CM | 11 |
| COREAD | 12 |
| ESCA | 3 |
| GBM | 4 |
| HC | 5 |
| HNSC | 7 |
| LGG | 1 |
| LUAD | 9 |
| LUSC | 5 |
| OV | 3 |
| PRAD | 3 |
| RCCC | 7 |
| SCLC | 4 |
| STAD | 5 |
| THCA | 7 |
| ICC | 11 |

Table 3: The list of the predicted driver-genes for each cancer type.

| Cancer type | Gene name | Number of mutations | Gene length | Mutation_Frequency |
|---|---|---|---|---|
| COREAD | MAPK1 | 33 | 98652 | 2989.454545 |
| HNSC | RRAGD | 18 | 47661 | 2647.83333 |
| CM/LUSC | PRKACB | 32 | 40800 | 1275 |
| COREAD | STIL | 61 | 64009 | 1049.327869 |
| UCEC | PKHD1 | 358 | 366777 | 1024.51676 |
| UCEC | ALDH1A1 | 53 | 52656 | 993.509434 |
| BLCA | SLC15A1 | 74 | 68875 | 930.7432432 |
| BRCA | TCF15 | 7 | 6274 | 896.2857143 |
| CM | RBP1 | 15 | 13425 | 895 |
| CM | SCD | 20 | 17817 | 890.85 |
| HC | PEBP1 | 11 | 9705 | 882.2727273 |
| LUSC | MAPK6 | 55 | 47052 | 855.4909091 |
| UCEC | SLC9A2 | 110 | 91817 | 834.7 |
| LUAD | PLOD1 | 52 | 40876 | 786.0769231 |
| STAD | ETNK1 | 28 | 19274 | 688.357143 |
| HNSC | TGM3 | 71 | 45113 | 635.3943662 |
| SCLC | SLC12A3 | 82 | 50644 | 617.6097561 |
| LGG | SULT2B1 | 39 | 23692 | 607.4871795 |
| GBM/CM | SULT1E1 | 33 | 18941 | 573.969697 |
| BRCA | PRPF19 | 30 | 16042 | 534.7333333 |
| HC | S100A11 | 9 | 4530 | 503.3333333 |
| GBM | PEX5 | 42 | 21122 | 502.9047619 |
| CM | TAL1 | 32 | 15426 | 482.0625 |
| CM | SUV39H1 | 26 | 12283 | 472.4230769 |
| LUAD | PDK4 | 28 | 13117 | 468.4642857 |
| HNSC | TFAP2B | 63 | 28888 | 458.5396825 |
| STAD | PGM3 | 60 | 27164 | 452.7333333 |
| HNSC | OSGIN1 | 29 | 13111 | 452.103448 |
| HC | SNRPB | 21 | 9219 | 439 |
| UCEC | CHST6 | 50 | 21905 | 438.1 |
| COREAD | CERCAM | 39 | 16872 | 432.615385 |
| BRCA | SCNN1G | 79 | 34161 | 432.4177215 |
| AML | RBM3 | 16 | 6813 | 425.8125 |
| COREAD | SLC28A2 | 57 | 23705 | 415.877193 |
| RCCC | ALG12 | 37 | 15253 | 412.243243 |
| UCEC | PITX3 | 28 | 11286 | 403.0714286 |
| COREAD | PPM1G | 74 | 28485 | 384.9324324 |
| CM/PRAD | PFKM | 62 | 23758 | 383.1935484 |
| LUSC/UCEC | RIT1 | 35 | 13108 | 374.5142857 |
| STAD | RUNX1T1 | 168 | 62714 | 373.297619 |
| COREAD | OBSCN | 412 | 153121 | 371.652913 |
| COREAD | SFTPB | 31 | 11425 | 368.5483871 |
| BRCA | POU3F1 | 8 | 2928 | 366 |
| BRCA | CXCL12 | 23 | 8036 | 349.3913043 |
| SCLC | COX17 | 24 | 7872 | 328 |
| COREAD | ZKSCAN2 | 66 | 21534 | 326.272727 |
| UCEC | SOX11 | 27 | 8719 | 322.9259259 |
| THCA | PIP | 24 | 7661 | 319.2083333 |
| PRAD/STAD/LUAD | PPP1R10 | 57 | 16907 | 296.6140351 |
| STAD | SRY | 3 | 887 | 295.6666667 |
| OV | POLR2A | 106 | 30238 | 285.2641509 |
| COREAD | SNRPD2 | 16 | 4539 | 283.6875 |
| RCCC | XCL1 | 20 | 5605 | 280.25 |
| LUSC | SLC9A5 | 85 | 23240 | 273.411765 |
| LUAD | PIM1 | 21 | 5283 | 251.5714286 |
| BLCA/RCCC | SFRP2 | 34 | 8487 | 249.6176471 |
| CM | STC1 | 53 | 12887 | 243.1509434 |
| CM | PYCR1 | 20 | 4854 | 242.7 |
| LUAD | ITPKA | 40 | 9702 | 242.55 |
| OV | POU3F3 | 19 | 4506 | 237.1578947 |
| STAD | SRMS | 32 | 7581 | 236.90625 |
| HNSC | TNKS1BP1 | 110 | 25311 | 230.1 |
| THCA | PIGR | 78 | 17945 | 230.0641026 |
| BRCA | SLC52A3 | 37 | 8505 | 229.864865 |
| BLCA | PLXNB1 | 112 | 25612 | 228.6785714 |
| BRCA | HAP1 | 54 | 12008 | 222.3703704 |
| COREAD | BRD8 | 101 | 21786 | 215.70297 |
| ESCA/SCLC | CCL3 | 9 | 1904 | 211.5555556 |
| HC | ANG | 26 | 5414 | 208.2307692 |
| BRCA | PROP1 | 20 | 4008 | 200.4 |
| HC | PGLYRP2 | 56 | 10859 | 193.9107143 |
| THCA | NOLC1 | 62 | 11696 | 188.6451613 |
| CM | LEFTY2 | 26 | 4895 | 188.2692308 |
| CLL | PTMS | 25 | 4636 | 185.44 |
| HNSC | CCNA1 | 62 | 10611 | 171.1451613 |
| LUAD | SIX3 | 26 | 4180 | 160.7692308 |
| CM | SRF | 59 | 9117 | 154.5254237 |
| COREAD | FES | 70 | 10731 | 153.3 |
| CLL | MAP2K7 | 73 | 10704 | 146.630137 |
| RCCC | GRB7 | 51 | 7319 | 143.5098039 |
| AML | SPTA1 | 543 | 76010 | 139.9815838 |
| BRCA | CCL1 | 21 | 2906 | 138.3809524 |
| THCA | RXRG | 64 | 8428 | 131.6875 |
| PA | FCER1G | 32 | 3952 | 123.5 |
| UCEC | PCOLCE | 51 | 5917 | 116.019608 |
| PRAD | MBD1 | 94 | 10733 | 114.1808511 |
| BRCA/HNSC | TAPBP | 104 | 10743 | 103.2980769 |
| LUAD | KRT18 | 38 | 3843 | 101.1315789 |
| UCEC | MYH4 | 294 | 26269 | 89.35034014 |
| LUSC | SSTR5 | 30 | 2674 | 89.13333333 |
| RCCC | SPHK1 | 34 | 2653 | 78.02941176 |
| GBM | NES | 113 | 8634 | 76.40707965 |
| ESCA | SSTR1 | 67 | 5065 | 75.59701493 |
| RCCC | PBX2 | 74 | 5451 | 73.66216216 |
| LUAD | PLP2 | 47 | 3285 | 69.89361702 |
| ESCA | RAG1 | 180 | 11748 | 65.26666667 |
| COREAD | S100A8 | 15 | 945 | 63 |
| UCEC | PNMT | 34 | 2023 | 59.5 |
| RCCC | FTL | 29 | 1571 | 54.17241379 |
| SCLC | GLMP | 57 | 3003 | 52.68421053 |
| THCA | IFNA8 | 20 | 1039 | 51.95 |
| BRCA | SLC1A3 | 58 | 2943 | 50.74137931 |
| LUAD | SOX15 | 40 | 1991 | 49.775 |
| THCA | EPHA10 | 100 | 4880 | 48.8 |
| UCEC | PTPRCAP | 53 | 2173 | 41 |
| BRCA | DAXX | 109 | 4458 | 40.89908257 |
| OV | PRM1 | 13 | 500 | 38.46153846 |
| BRCA | FKBPL | 45 | 1584 | 35.2 |
| THCA | SOD1 | 332 | 9309 | 28.03915663 |
| GBM | SPEG | 224 | 6051 | 27.01339286 |
| BLCA | PGK2 | 96 | 1690 | 17.60416667 |
| BRCA | HIST1H2AC | 47 | 546 | 11.61702128 |

Table 4: List of the rare mutations within the predicted head and neck squamous cell carcinoma driver genes. The mutations are filtered by minor allele frequency (MAF) in ExAC with <0.01

| Gene Name | Mutations | Number of patients share this mutation | ExAC_Freq | SIFT score | Polyphen2 score | MutationTaster score | MutationAssessor score | FATHMM score | VEST3 score | Unscaled/Raw CADD | PHRED-scaled CADD | Type | Prediction |
|---|---|---|---|---|---|---|---|---|---|---|---|---|---|
| TGM3 | rs114998364 | 1 | 0.0027 | 0.844 | 0.001 | 1 | 0.345 | -0.16 | 0.066 | 0.429 | 6.863 | heterozygous | **tolerated** |
| TGM3 | rs200294064 | 1 | $0.6 \times 10^{-4}$ | 0.063 | 0.936 | 1 | 1.965 | 1.52 | 0.618 | 4.825 | 24.8 | heterozygous | **deleterious** |
| TGM3 | rs539884508 | 1 | 0.0004 | 0.595 | 0.002 | 1 | 0.55 | -0.72 | 0.046 | -1.38 | 0.004 | heterozygous | **tolerated** |
| OSGIN1 | rs138964906 | 1 | 0.0007 | 0.83 | 0.038 | 0.997 | 0.9 | 1.34 | 0.05 | 1.214 | 11.82 | heterozygous | **tolerated** |
| TNKS1BP1 | rs747045370 | 1 | 0 | 0.269 | 0.042 | 1 | 0.345 | 1.61 | 0.106 | 0.415 | 6.743 | heterozygous | **tolerated** |
| TNKS1BP1 | rs115694714 | 3 | 0.0030 | 0.007 | 0.993 | 0.626 | 0.695 | 0.98 | 0.463 | 4.699 | 24.6 | heterozygous | **deleterious** |
| TNKS1BP1 | rs79359831 | 2 | 0.0032 | 0.121 | 0.017 | 1 | 1.61 | 0.97 | 0.189 | 1.967 | 16 | heterozygous | **deleterious** |
| TNKS1BP1 | rs748159115 | 1 | 0 | 0.535 | 0.025 | 1 | 0.41 | 1.45 | 0.136 | 0.7 | 8.826 | heterozygous | **tolerated** |
| CCNA1 | rs113565588 | 3 | 0.0002 | 0.306 | 0.001 | 1 | 0 | 2.39 | 0.151 | -0.001 | 2.574 | heterozygous | **tolerated** |
| TAPBP | rs145837211 | 3 | 0.0052 | 0 | 0.002 | 1 | . | 1.32 | 0.101 | 3.795 | 23.4 | heterozygous | cannot assess |

Table 5: List of the rare mutations within the predicted colorectal driver genes. The mutations are filtered by minor allele frequency (MAF) in ExAC with <0.01

| Gene name | Mutation | Number of patients share this mutation | ExAC_Freq | PolyPhen2 score | SIFT score | MutationAssessor score | MutationTaster score | FATHMM score | VEST3 score | Unscaled/Raw CADD | PHRED-scaled CADD | Type | Prediction |
|---|---|---|---|---|---|---|---|---|---|---|---|---|---|
| FES | rs761486582 | 2 | 0 | 0.988 | 0.003 | 1.205 | 1 | -1.67 | 0.532 | 3.615040 | 25.5 | heterozygous | **deleterious** |
| OBSCN | rs377742814 | 2 | $0.3 \times 10^{-4}$ | 0.989 | 0.03 | 2.61 | 1 | -0.27 | 0.588 | 2.928777 | 23.5 | heterozygous | **deleterious** |
| OBSCN | rs201725133 | 2 | $0.8 \times 10^{-4}$ | . | . | . | 1 | . | . | 6.609900 | 36 | heterozygous | cannot assess |
| OBSCN | rs775164465 | 2 | 0 | 0.001 | 0.525 | 0.02 | 1 | 0.62 | 0.131 | -0.369758 | 0.224 | heterozygous | **deleterious** |
| OBSCN | rs372037309 | 2 | $0.5 \times 10^{-4}$ | 0.737 | 0 | 1.78 | 1 | 1.07 | 0.388 | 3.478409 | 25 | heterozygous | **deleterious** |
| OBSCN | rs373638525 | 2 | 0.00012 | 0.989 | 0.022 | 0.885 | 0.867 | -0.21 | 0.317 | 3.502569 | 25.1 | heterozygous | **deleterious** |
| OBSCN | rs770745639 | 2 | 0 | 0.023 | 0.026 | 0.95 | 0.999 | -0.01 | 0.062 | 3.252189 | 24.3 | heterozygous | **deleterious** |
| OBSCN | rs776318810 | 2 | 0 | 0.029 | 0.218 | 1.505 | 1 | -0.23 | 0.186 | 0.434124 | 8.744 | heterozygous | **tolerated** |
| OBSCN | rs749249863 | 2 | 0 | 0.877 | 0.215 | 0.655 | 1 | -0.28 | 0.196 | 2.196906 | 21.3 | heterozygous | **tolerated** |
| OBSCN | rs188024384 | 2 | 0.0015 | 0.105 | 0.139 | 0.69 | 1 | -0.21 | 0.258 | 0.680169 | 10.93 | heterozygous | **deleterious** |
| OBSCN | rs756875131 | 2 | 0 | 0.904 | 0.085 | 2.385 | 1 | -0.35 | 0.177 | -0.091695 | 1.480 | heterozygous | **deleterious** |
| OBSCN | rs201854668 | 2 | 0.00124 | 0.999 | 0.088 | 2.045 | 1 | -0.2 | 0.416 | 7.344976 | 38 | heterozygous | **deleterious** |
| OBSCN | rs188531451 | 2 | $0.8 \times 10^{-4}$ | 0.974 | 0.051 | 2.465 | 1 | 0.29 | 0.31 | 3.299496 | 24.4 | heterozygous | **deleterious** |
| OBSCN | rs559123841 | 2 | $0.8 \times 10^{-4}$ | . | 0.687 | . | 1 | 3.72 | 0.025 | -0.020337 | 2.326 | heterozygous | cannot assess |
| OBSCN | rs62621832 | 2 | 0.00709 | 1 | 0 | 3.14 | 1 | 2.1 | 0.664 | 3.530120 | 25.2 | homozygous | **deleterious** |
| OBSCN | rs199501426 | 2 | 0.00022 | 0.156 | 0.071 | 2.305 | 1 | -0.3 | 0.499 | 2.536169 | 22.7 | heterozygous | **deleterious** |

Table 6: List of overlapping predicted driver genes with the current COSMIC confirmed cancer driver genes

| Gene Name |
|---|
| MAPK1 |
| STIL |
| ETNK1 |
| TAL1 |
| RUNX1T1 |
| PIM1 |
| FES |
| DAXX |

Table 7: The comparison to ICGC Data with the somatic variants detected by us in nasopharyngeal carcinoma and colorectal adenocarcinoma.

| dpSNP ID | Mutation ID | Functional Impact | Tumor Type | Gene Name |
|---|---|---|---|---|
| rs146355154 | _ | _ | _ | TGM3 |
| rs200294064 | MU4596969 | Low | Biliary Tract/Esophageal cancer | TGM3 |
| rs539884508 | _ | _ | _ | TGM3 |
| rs138964906 | _ | _ | _ | OSGIN1 |
| rs747045370 | _ | _ | _ | TNKS1BP1 |
| rs115694714 | _ | _ | _ | TNKS1BP1 |
| rs79359831 | MU80430 | Low | Colon cancer | TNKS1BP1 |
| rs748159115 | MU4392892 | Low | Skin cancer | TNKS1BP1 |
| rs113565588 | MU113413488 | Low | Lung cancer | CCNA1 |
| rs145837211 | _ | _ | _ | TAPBP |
| rs761486582 | MU1988312 | High | Endometrial cancer | FES |
| rs377742814 | _ | _ | _ | OBSCN |
| rs201725133 | _ | _ | _ | OBSCN |
| rs775164465 | _ | _ | _ | OBSCN |
| rs372037309 | _ | _ | _ | OBSCN |
| rs373638525 | _ | _ | _ | OBSCN |
| rs770745639 | MU4461516 | Low | Skin cancer | OBSCN |
| rs776318810 | MU4725242 | Low | Gastric cancer | OBSCN |
| rs749249863 | MU67306141 | Low | Breast/Pediatric Brain cancer | OBSCN |
| rs188024384 | MU60415801 | Low | Skin cancer | OBSCN |
| rs756875131 | MU4724704 | Low | Gastric cancer | OBSCN |
| rs201854668 | MU4588766 | Low | Head and Neck cancer | OBSCN |
| rs188531451 | _ | _ | _ | OBSCN |
| rs559123841 | _ | _ | _ | OBSCN |
| rs62621832 | _ | _ | _ | OBSCN |
| rs199501426 | MU70565483 | Low | Esophageal/Gastric cancer | OBSCN |

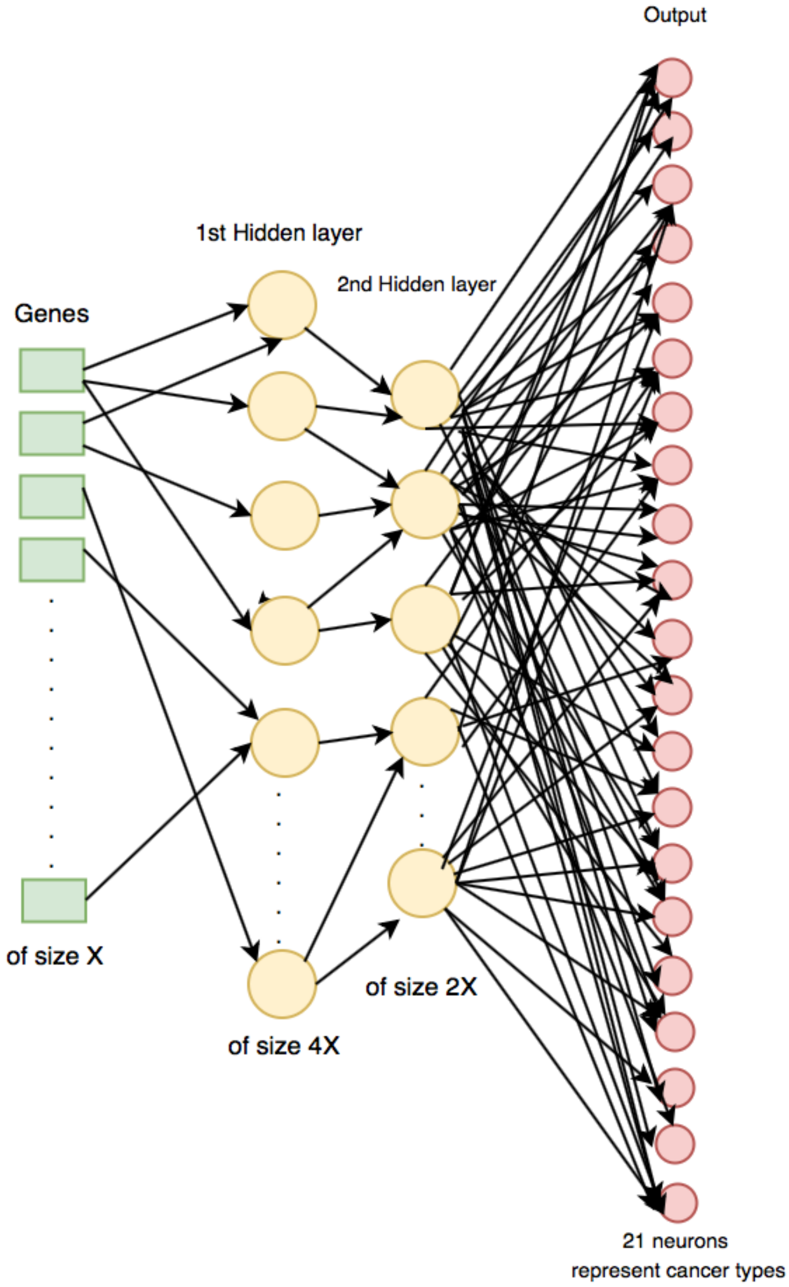


Figure 1: Neural network architecture of number of genes as a vector representation of size X and two hidden layers of size 4X and 2X and the output of different 21 neurons for each 20 cancer types plus not identified gene class.
